# A gene essentiality signature enables predicting the mechanism of action of drugs

**DOI:** 10.1101/2022.11.07.514541

**Authors:** Wenyu Wang, Jie Bao, Shuyu Zheng, Shan Huang, Jehad Aldahdooh, Yinyin Wang, Johanna Eriksson, Ziaurrehman Tanoli, Xuepei Zhang, Massimiliano Gaetani, Jing Tang

**Affiliations:** Research Program in Systems Oncology, Faculty of Medicine, University of Helsinki, Helsinki 00290, Finland; Department of Radiation Oncology, Second Affiliated Hospital, Xi’an Jiaotong University, No.157, Xi Wu Road, Xi’an, 710004, Shaanxi, China; Chemical Proteomics Core Facility, Division of Chemistry I Department of Medical Biochemistry and Biophysics, Karolinska Institutet, Stockholm, Sweden; Chemical Proteomics Unit, Science for Life Laboratory (SciLifeLab), Stockholm, Sweden; Chemical Proteomics, Swedish National Infrastructure for Biological Mass Spectrometry (BioMS), Stockholm, Sweden

## Abstract

Cancer drugs often kill cells independent of their putative targets, suggesting the limitation of existing knowledge on the mechanisms of action. In this study, we explored whether the integration of loss-of-function genetic and drug sensitivity screening data can define a gene essentiality signature to better understand the drug target interactions. We showed that our gene essentiality signature can predict drug targets more accurately than chemical fingerprints and drug-perturbated gene expression signatures. We further showed how gene essentiality signature can help identify mechanisms of action of drugs *de novo*, including the EGFR inhibitor lapatinib, and drugs associated with DNA mismatch repair. Finally, we established gene essentiality signatures for noncancer drugs and used them to predict their anticancer targets. We have successfully validated the target predictions for multiple noncancer drugs, using cell-based drug target deconvolution by the proteome integral solubility alteration assay. Our study provides a novel signature of drugs that may facilitate the rational design of drug repurposing.

## Introduction

Drug-target interactions trigger a cascade of downstream molecular modulations, leading to phenotypic changes of cancer cells such as viability and proliferation. Although many candidate drugs are tested in clinical trials, the full knowledge of their mechanisms of action is often lacking. For example, a recent study has shown that drugs kill cancer cells even when their putative targets are depleted, suggesting the limitations of existing drug target information^1^. The incomplete drug target information also prevents biomarker identification and, in the end, may lead to treatment failure due to inaccurate patient stratification^2^.

Drug discovery process mainly falls into two categories, target-based or phenotype-based^3^. The target-based drug discovery starts with a particular target gene or a disease pathway, followed by the identification of potent chemical hits^4,5^. In comparison, phenotype-based drug discovery select candidate compounds that lead to desired cellular phenotypes, which are then forwarded for target deconvolution^6^. Direct target deconvolution approaches, *e.g*., cell-free affinity experiments, are typically conducted in a hypothesis-driven manner with a handful purified candidate proteins^3^. Advanced target deconvolution assays, such as Thermal Proteome Profiling (TPP)^7^ and Proteome Integral Solubility Alteration (PISA)^8^, allow interrogating proteome-wide drug-target interactions using cell lysate or intact cells. TPP is a time- and effort-consuming target deconvolution technique, where a full multiplex experiment (batch) can be used for one or maximum two biological replicates and testing one molecule versus control in duplicates already requires a set of several multiplex proteomics experiment. The PISA assay provided a revolution in throughput, robustness, missing values, number of biological replicates and statistical power, and multiplicity of different compounds/conditions, up to five in triplicates compared to controls per batch. In its new 18-plex format^9^, PISA assay enables a more sustainable proteomics-based target deconvolution, for example with 5 drugs or conditions in triplicate, plus the relative control, in one assay, but this still need further developments to enable sustainability of very large datasets. For example, a recent phenotype-based drug screening study showed that many approved noncancer drugs can efficiently kill cancer cells^10^. Despite the initial evidence, why and how these drugs affected cancer cells remains poorly understood, partly due to the limited capacity of hypothesis-free target deconvolution approaches. To our knowledge, an unbiased proteome-wide drug target deconvolution on a large panel of cell lines is currently unavailable^11^. Therefore, the need to develop systems medicine approaches to understand the mechanisms of actions is strong, especially for the repurposing of noncancer drugs.

One common strategy to study the mechanisms of action of drugs is gene expression signature analysis. For example, the L1000 assay^12^ or the PLATE-Seq assay^13^ allows measuring drug-induced transcriptomics changes. The gene expression signatures greatly enhanced the identification of drug targets and their downstream effectors, providing critical information to understand drug mechanisms at the pathway level^14–17^. Despite the success of many machine learning models in exploiting the gene expression signatures, it should be noted that the gene expression signature data has limited coverage. Notably, the commonly used LINCS-L1000 dataset includes only 978 consensus genes for dozens of cell lines. PANACEA is a more recent study expected to increase the coverage of gene expressions, but with a limited tumor context consisting of 25 cell lines^15^.

On the other hand, loss-of-function genetic screens have been applied to study genetic dependencies in cancer, termed gene essentiality profiles^18^. In contrast to the drug-induced gene expression assays, the gene essentiality screening allows rapid genetic perturbations for a large number of genes and cell lines, thanks to the pooled CRISPR and RNAi techniques^19–22^. For example, the dependency mapping (DepMap) study has profiled the gene essentiality profiles of more than 17k genes for a panel of 1070 cancer cell lines. Meanwhile, the same panel of cell lines has been tested in drug sensitivity screening (i.e., CTRP and GDSC), which forms the rationale to predict drug sensitivities with gene essentiality profiles^10,23^. However, there exist few studies on leveraging the gene essentiality profiles for drug target prediction. Our study was motivated by Gonçalves et. al., which reported interesting overlaps between drug sensitivity associated essential genes and their putative targets^24^. Despite the initial evidence, the gene essentiality features that were predictive for drug-target identification were not established. Furthermore, there is a lack of systematic comparison of the gene essentiality-based features against conventional drug features, such as gene expression signatures, or structure-based fingerprints. Thirdly, it was unclear whether the gene-essentiality-based drug target prediction approaches can be generalized for noncancer drugs.

In this study, we aimed to solve abovementioned limitations by deriving a novel gene essentiality signature for a drug. Our study was based on the complementarity of drug sensitivity and gene essentiality screens. Despite targeting molecules using different approaches, both screens assess the phenotypic effects of perturbations (**Figure 1A**). We developed a regression model to deconvolute drug sensitivity into its gene-level perturbation effects by leveraging CRISPR- and RNAi-based functional genetics data. The estimated model coefficients were determined as the gene essentiality signatures of the drugs. We evaluated the gene essentiality signatures as compared to gene expression signatures and chemical fingerprints, in both drug target prediction and mechanisms of action discovery (**Figure 1B-C**). Finally, we applied the gene essentiality signatures for noncancer drugs and determined their potential targets that may explain their anti-cancer efficacies (**Figure 1D**).

**Figure 1.**
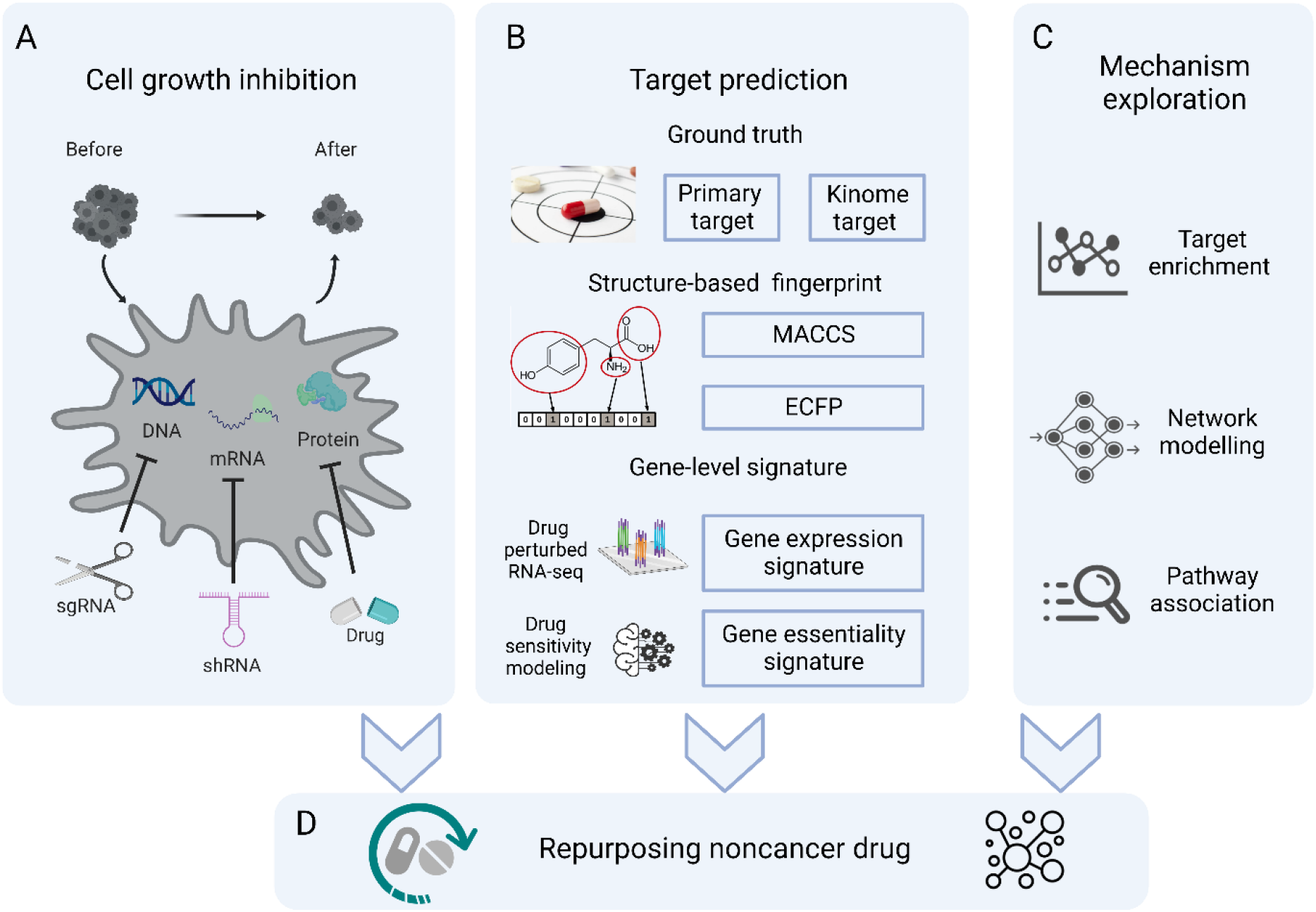
Schematic of the study design. A. Cell growth inhibition at different levels of perturbations enables the use of gene essentiality features to predict drug sensitivity. sgRNA: single guide RNA; shRNA: short hairpin RNA. B. The gene essentiality signatures are compared with chemical fingerprints and gene expression signatures for drug-target prediction. MACCS: Molecular ACCess Systems keys fingerprint; ECFP: Extended Connectivity Fingerprint. C. The gene essentiality signatures can also be used for *de novo* exploration of mechanisms of action, including target enrichment, network modelling, and pathway association. D. Application of the gene essentiality signatures to repurpose non-cancer drugs.

## Results

### Establishing the gene essentiality signatures of drugs

For each drug, we established a regression model between drug sensitivity and gene essentiality of the cell lines, based on the rationale that chemical or genetic perturbations targeting the same cancer signaling pathways lead to similar growth inhibition effects (**Figure 2A**). To minimize the model complexity due to data integration noise, we employed an L1 penalized ridge regression to estimate the coefficients for all the gene essentiality features. We first evaluated whether the regression models built on the gene essentiality features were generally predictive of drug sensitivity. More specifically, we considered three commonly used variants of gene essentiality features that were determined using different techniques, namely CERES, DEMETER2, and CES (**Methods section**). We showed the model performance for the CTRP and GDSC data separately.

**Figure 2.**
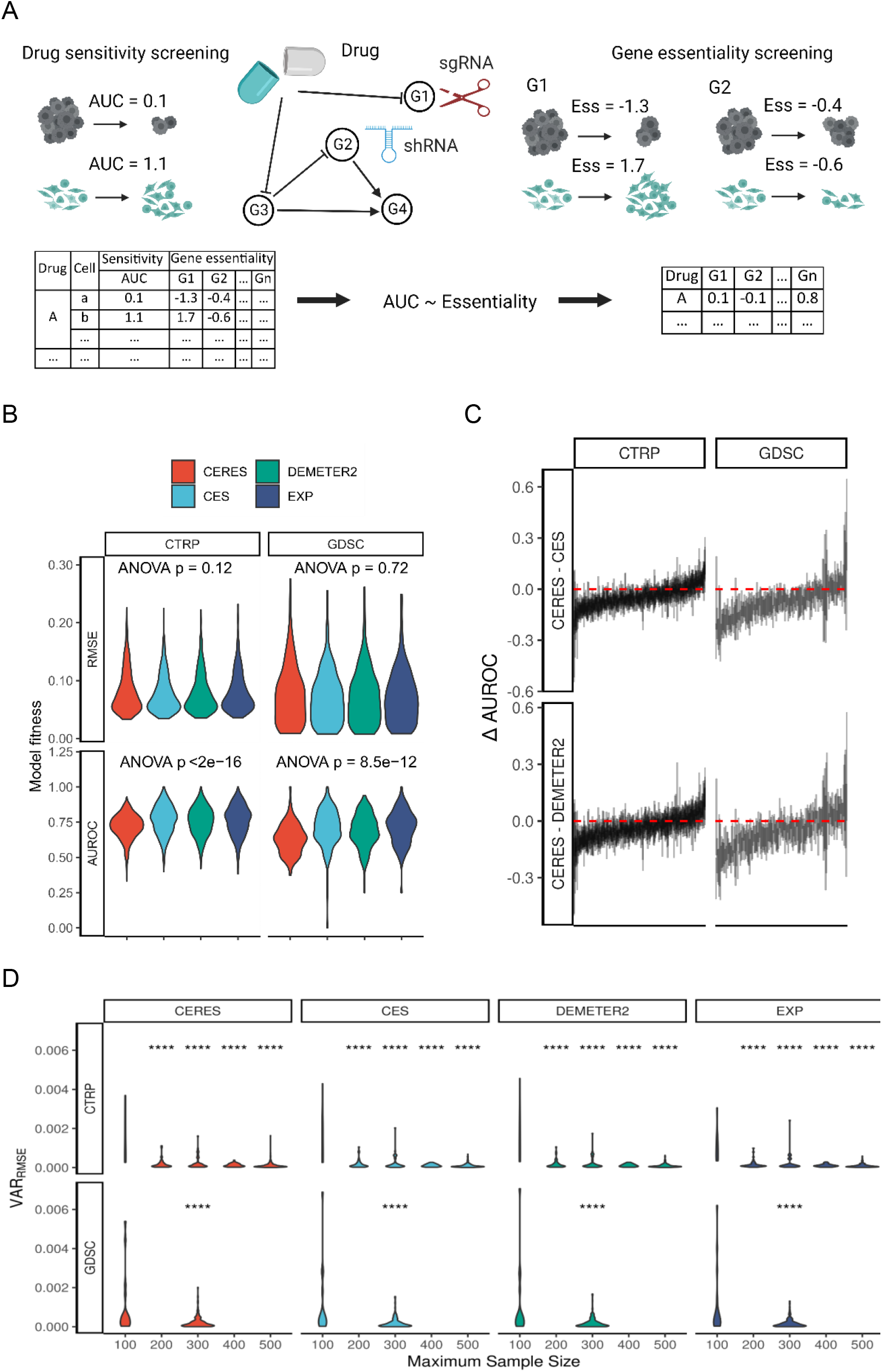
Establishing gene essentiality signatures of drugs. A. Rationale for modeling of drug sensitivity with gene essentiality profiles. In drug sensitivity screening, a drug induces cell growth inhibition by triggering a cascade of downstream signaling events. In gene essentiality screening, cell growth inhibition is achieved by either shRNA-mediated knocking down or sgRNA-mediated knocking out. For each drug, a drug sensitivity regression model across multiple cell lines can be built by treating their gene essentiality profiles as features. B. Model fitness of gene essentiality features, including CERES, DEMETER2, and CES, when comparing to gene expression features (EXP). C. Differences in AUROC between CERES, DEMETER2, and CES. Each vertical line along the x axis represents the 95% confidence interval of the difference for a drug. Red horizontal lines show the reference of zero difference. D. Variance of RMSE for each feature type grouped by drugs with different sample sizes. ****: p < 0.0001 by Dunnett’s test using maximum sample size of 100 as the reference.

As shown in **Figure 2B and Supplementary Figure 1**, all the three versions of gene essentiality features yield comparable model fitness (defined as the spearman correlation between model prediction and ground truth of drug sensitivity values), to the same level of using the gene expression feature, which is known to be predictive for drug sensitivity^25,26^. The RMSE did not show significant differences among the three gene essentiality features (one-way ANOVA test P-value = 0.12 for CTRP and GDSC). For the CERES signature, we observed a slight decrease in AUROC compared to the gene expression feature (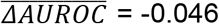, Tukey’s test P-value = 6.8×10^-11^ for CTRP, and 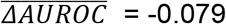, Tukey’s test P-value < 2×10^-16^ for GDSC), while the CES and DEMETER2 gene essentiality features were equally predictable (all the Tukey’s test P-values > 0.05, **Supplementary Table 1**). Furthermore, CERES and CES showed no difference when predicting 91.6% of the CTRP drugs and 91.4% of the GDSC drugs, while CERES and DEMETER2 showed no difference when predicting 95.4% of the CTRP and 95.5% of the GDSC drugs (**Figure 2C and Supplementary Tables 2-3**). Therefore, we considered that all the three types of gene essentiality features have similar performance, and therefore took the average of their gene essentiality coefficients to derive a consensus gene essentiality signature for each drug, consisting of 10,624 genes.

On the other hand, we found that the drugs with smaller sample sizes (i.e., number of cell lines) tend to yield a larger variance of RMSE in the cross-validation (**Figure 2D**), especially if sample size is less than 100 (all Dunnett test P-value < 0.001, **Supplementary Table 4**). To ensure the reliability of the gene essentiality signatures, we considered only the drugs tested with more than 100 cell lines for the subsequent analyses.

### The gene essentiality signatures predict targets of cancer drugs

We explored how the gene essentiality signatures can predict drug targets in a supervised setting (**Figure 1B**). We employed a similarity-based method that showed superior performance in a recent DREAM Challenge competition for drug target predictions (**Methods section**)^15^. The performance of the gene essentiality signature was compared with that of the gene expression signatures, which are the transcriptomics changes after drug treatment. We also evaluated the MACCS and ECFP fingerprints generated using the chemical structures of the drugs.

We observed that the gene essentiality signature achieved top performance across CTRP and GDSC datasets (**Figure 3A and Supplementary Table 5**). For example, for the GDSC dataset, when using the primary targets of drugs as the ground truth, the gene essentiality signature achieved an average AUC of 0.61, as compared to 0.55, 0.47 and 0.43 for the gene expression signature, ECFP and MACCS fingerprints, respectively (**Figure 3A, upper panel**). In addition, we also examined the performance in predicting a broader spectrum of secondary targets. We considered the kinome-wide drug target interaction scores as the ground truth and examined the prediction performance by binarizing the scores with a threshold of 0.4. We observed a similar result for the CTRP and GDSC datasets, where the gene essentiality signatures generally outperformed the other drug signatures (**Figure 3A, lower panel**).

**Figure 3.**
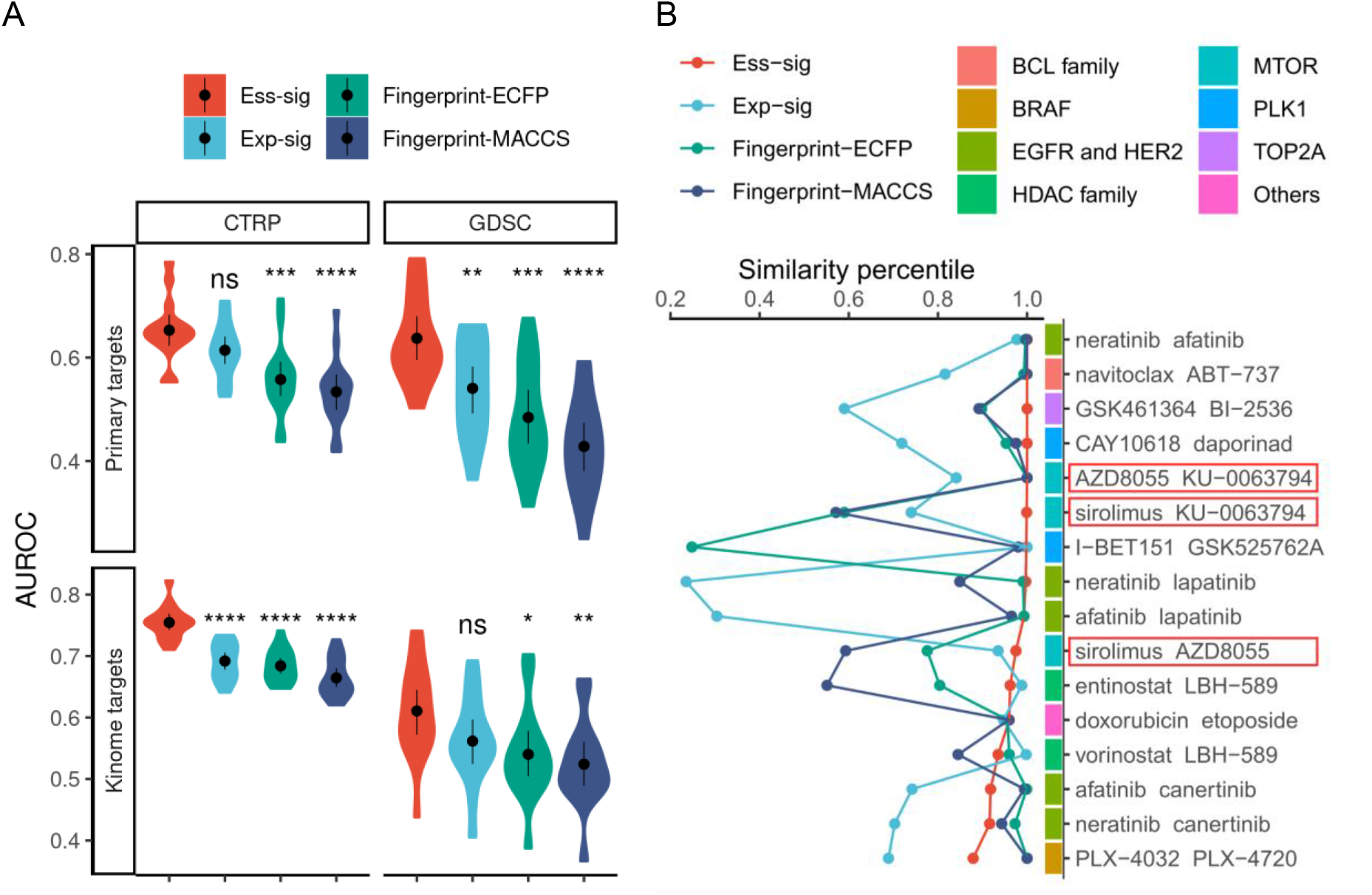
Gene essentiality signatures improve drug target prediction. A. Accuracy of drug target prediction for CTRP and GDSC datasets. ns: p > 0.05, *: p <= 0.05, **: p <= 0.01, ***: p <= 0.001, ****: p <= 0.0001 by Dunnett’s test with the gene essentiality signature as the reference. Ess-sig: gene essentiality signature, Exp-sig: gene expression signature. B. Ranking of the ground-truth repurposable drug pairs by their similarity using different signatures.

We are also interested in whether the gene essentiality signatures can prioritize drug pairs that share similar drug targets and drug sensitivity profiles. We reasoned that such drug pairs are top candidates for drug repurposing, as their drug sensitivity similarities are corroborated by the shared mechanisms of action. We determined such drug pairs as the ground truth, and then examined whether they can be prioritized by the gene essentiality signatures. As shown in **Supplementary Figure 2** and **Supplementary Table 6**, we found that the gene essentiality signatures successfully prioritized the ground truth drug pairs in both CTRP (median percentile = 93.4% for n = 150 drug pairs when considering the primary targets, and median percentile = 97.6% for n = 70 drug pairs when considering the kinome-wide targets), and GDSC datasets (median percentile = 90.8% for n = 29 dug pairs when considering the primary targets, and median similarity percentile = 95.4% for n = 23 drug pairs when considering kinome-wide targets).

The ranking of the ground truth drug pairs using different drug signatures is shown in **Figure 3B**. The gene essentiality signature was able to prioritize a majority of the ground truth drug pairs, for which the structure fingerprints and gene expression signatures failed to predict, indicating that the gene essentiality signature provides novel information to improve drug target prediction. For example, the gene essentiality feature was able to prioritize all the mTOR inhibitor drug pairs involving AZD8055, KU-0063794, and sirolimus (all with ranking higher than 97.5% percentile). In contrast, the chemical fingerprints were able to prioritize only the AZD8055-KU-0063794 pair, as they share the same pyridopyrimidine structure^27^ that is lacking in sirolimus. The gene expression signature prioritized the sirolimus-AZD8055 pair with a relative high rank at 93.5%, but not for sirolimus-KU-0063794 and AZD8055-KU- 0063794 pairs (rank at 74% and 84%, respectively).

### Gene essentiality signature facilitates exploration of mechanisms of action

After confirming the improved performance of gene essentiality signatures in the drug target prediction, we explored if a drug’s targets can be enriched in its gene essentiality signatures, despite that the gene essentiality signatures were determined without leveraging drug target information. Note that the gene essentiality signature is a vector of genes with feature importance values, so we asked whether the primary targets of drugs can be identified as top-ranking genes in the signature. As shown in **Figure 4A**, many drugs had their primary targets enriched in their top essentiality signature genes (Fisher’s test P = 1.2×10^-7^). For example, the primary targets for 30.4% of CTRP drugs and 57.1% of GDSC drugs were found within the top 50 genes in the gene essentiality signatures. In contrast, the primary targets showed no enrichment in the bottom genes of the gene essentiality signatures (Fisher’s test P > 0.05). This suggests that most drugs induced inhibitory effects on their targets, consistent with the inhibition effects induced by the gene essentiality screens. On the other hand, drug targets were much less enriched in the gene expression signatures (Fisher test P-value < 2.2×10^-16^). For example, only 3.7% and 3.1% of the CTRP drugs had their primary targets identified in the top 50 and bottom 50 genes, respectively. Likewise, using the gene essentiality signature to classify drug targets resulted in higher average AUC of 0.66 (CTRP) and 0.78 (GDSC), compared to an AUC of 0.47 (CTRP) and 0.53 (GDSC) for gene expression signatures (t-test P-value = 6.0×10^-5^ and 9.0×10^-4^ for CTRP and GDSC, respectively, **Figure 4B-4C and Supplementary Figure 4**).

**Figure 4.**
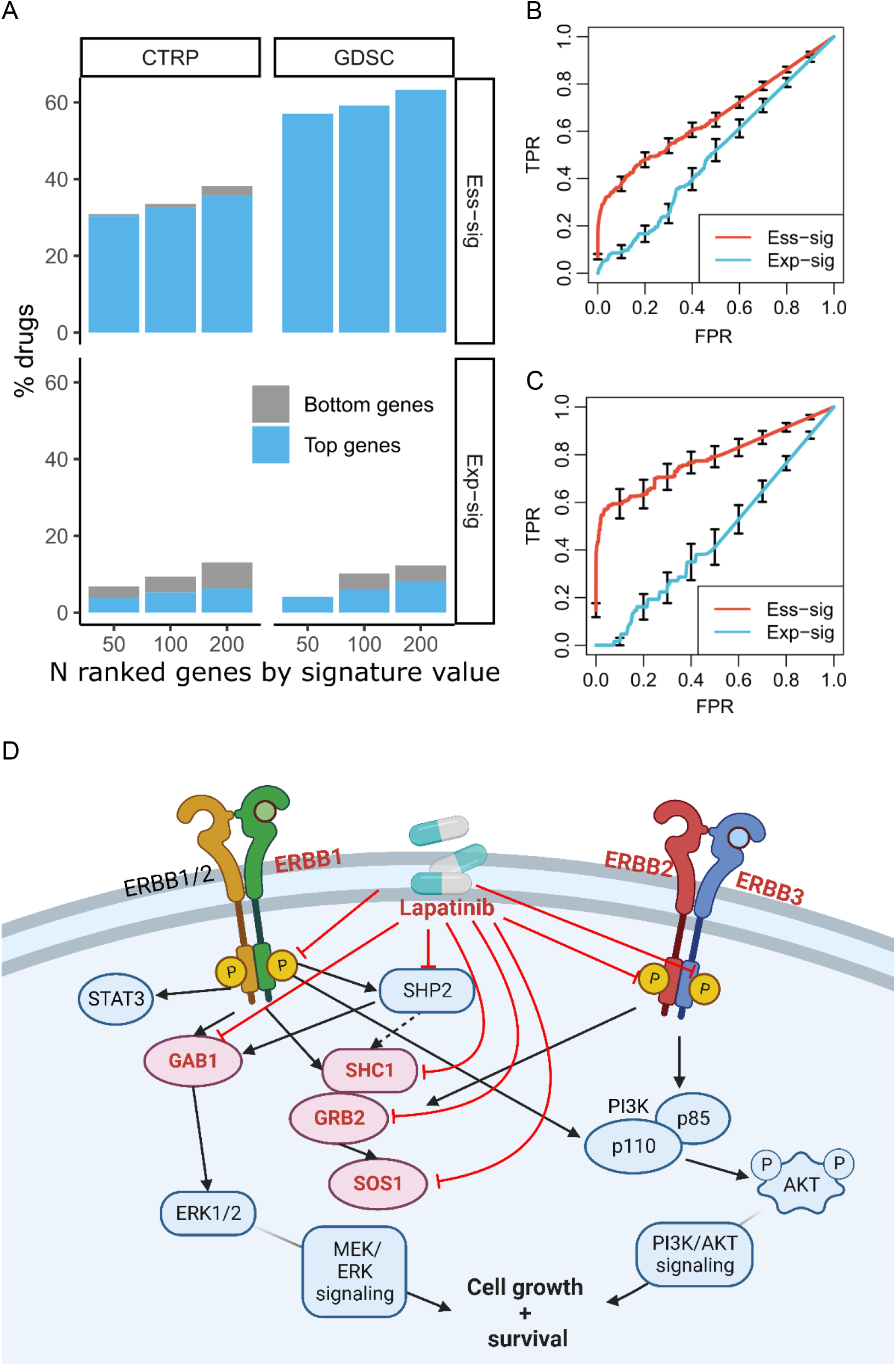
Mechanism of action analysis using the gene essentiality signature. A. Percentage of drugs with putative targets recovered by the top and bottom genes in their gene signatures. B-C. ROC curves of primary targets enrichment for CTRP and GDSC datasets, respectively. D. The mechanisms of action of lapatinib, with top genes in the gene essentiality signature of lapatinib highlighted.

We next checked whether the gene essentiality signatures may further capture the downstream pathways of the drugs. We found that the top and the bottom genes cover not only primary targets but also their neighboring genes in the PPI network (**Supplementary Figure 4**). For example, the top 50 and bottom 50 genes together can already identify the 1st-degree neighbors of targets for 33.5% of CTRP drugs and more than half of the GDSC drugs. This showed the gene essentiality signature’s ability to reveal both primary targets and downstream changes. In comparison, the top and bottom genes in the gene expression signatures identify mostly the 1st-degree neighbors (52.6%), suggesting that drugs often do not affect the expression of their target gene but only the downstream factors. We showed an EGFR inhibitor lapatinib as an example, for which the top 10 genes from the gene essentiality signature not only identified the primary targets, namely ERBB1(EGFR) and ERBB2 (HER2), but also revealed important downstream genes in the EGFR signaling pathway, including ERBB3^28^, GAB1^29,30^, and SOS1^30^ (**Figure 4D**).

To further map the gene essentiality signatures to biological processes, we implemented a gene set enrichment analysis to determine the pathways that are enriched in the gene essentiality signatures (**Supplementary Tables 7 - 8**). Here we highlighted the drugs for which the DNA mismatch repair pathway was enriched in their gene essentiality signatures. As shown in Table 1, many drugs are known to impact cell cycle and genome integrity, suggesting the validity of the gene essentiality signatures. DNA mismatch repair pathway may affect tumor mutation burden and immune cell infiltration, and thus is considered a potential target for enhancing immunotherapy^31,32^. Indeed, we found three approved drugs (talazoparib, cytarabine, and irinotecan) that are under investigation in combination with immunotherapy in recent clinical trials (**Supplementary Table 9**). For example, the PARP inhibitor talazoparib is combined with immune checkpoint inhibitors in multiple cancers, including urothelial carcinoma (NCT04678362), lung cancer (NCT04173507), as well as breast and ovarian cancer (NCT03330405). The remaining compounds enriched in the DNA mismatch repair pathway may also be worthy of further exploration as sensitizers for immune checkpoint inhibitors.

**Table 1.** List of drugs for which the DNA mismatch repair pathway was enriched in their gene essentiality signatures.

| DRUG NAME | P<sub>adj</sub> | ES | NES | REPORTED MECHANISM | DRUG TYPE |
| --- | --- | --- | --- | --- | --- |
| AZD7762 | 6.11×10<sup>-5</sup> | 0.75 | 2.40 | Cell cycle | Targeted |
| MK-8776 | 4.95×10<sup>-4</sup> | 0.71 | 2.21 | Cell cycle | Targeted |
| Wee1 Inhibitor | 1.24×10<sup>-3</sup> | 0.71 | 2.26 | Cell cycle | Targeted |
| MK-1775 | 2.97×10<sup>-3</sup> | 0.67 | 2.07 | Cell cycle | Targeted |
| Talazoparib | 3.40×10<sup>-3</sup> | 0.70 | 2.17 | Genome integrity | Targeted |
| AZD6738 | 8.53×10<sup>-3</sup> | 0.64 | 2.05 | Genome integrity | Targeted |
| VE-822 | 8.54×10<sup>-3</sup> | 0.66 | 2.04 | Genome integrity | Targeted |
| Cytarabine | 3.51×10<sup>-2</sup> | 0.61 | 1.96 | Other | Chemotherapeutic |
| Irinotecan | 4.18×10<sup>-2</sup> | 0.62 | 1.95 | DNA replication | Chemotherapeutic |

### Gene essentiality signatures help target prediction for the PRISM drugs

Profiling Relative Inhibition Simultaneously in Mixture (PRISM)^33^ is a recent technology that increases the throughput of drug sensitivity screening, with which researchers have discovered multiple noncancer drugs with efficacy in killing cancer cells^10^. Over half (53.4%) of the 1448 compounds probed in their multi-dose secondary screen were developed for indications other than cancer, mostly already approved^10^ (n = 406). However, these noncancer drugs, such as antibiotics and anti-parasite drugs, often lack relevant cancer target information (**Figure 5A and Supplementary Table 10**). Therefore, we tested whether the gene essentiality signature can be applied to the PRISM data to elucidate the mechanisms of action of these promising drugs for cancer treatment.

**Figure 5.**
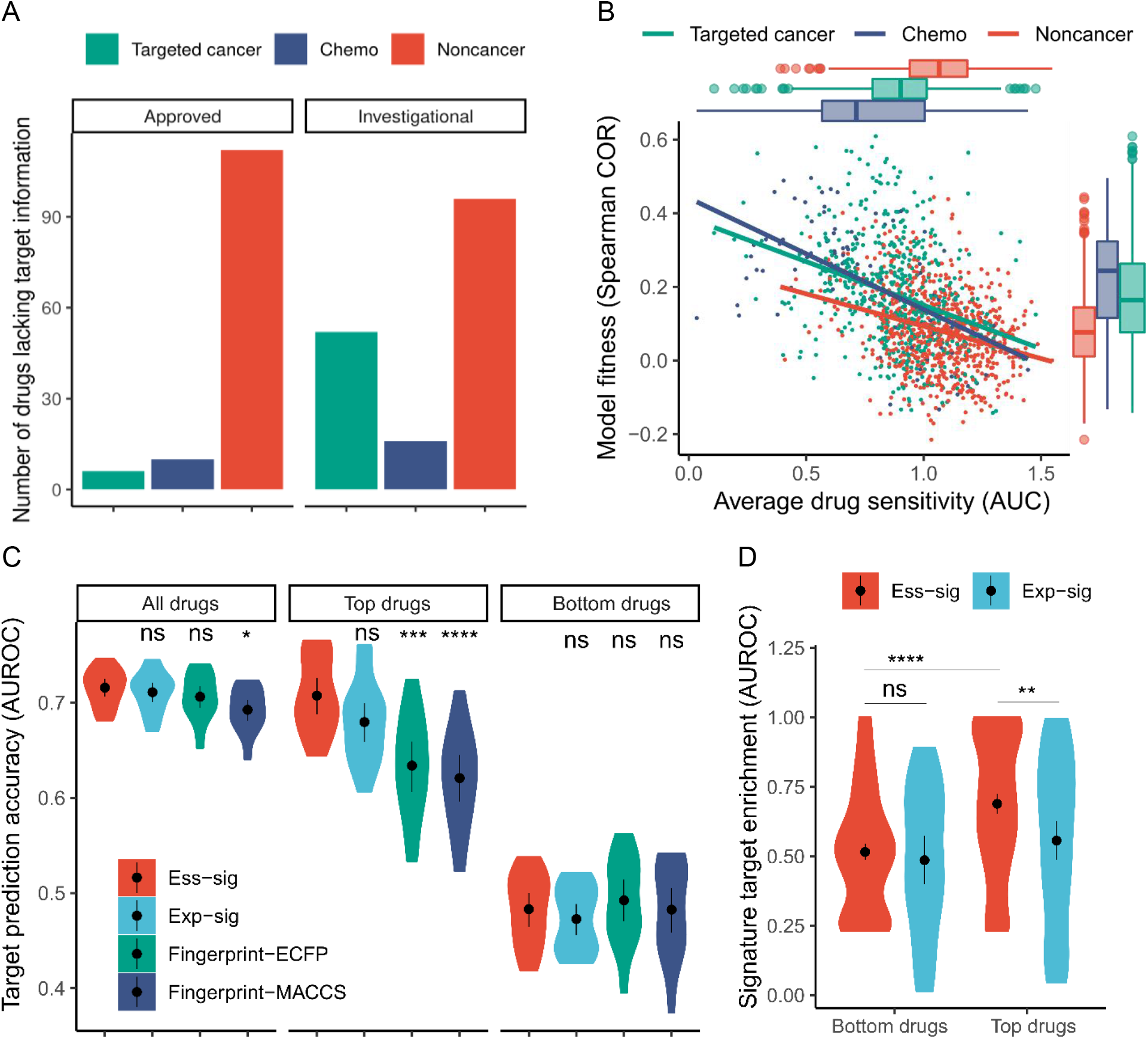
Application of gene essentiality signatures to predict targets of PRISM drugs. A. Numbers of drugs with missing target information in different categories B. Model fitness in association with the average drug sensitivity (AUC). Noncancer drugs tend to have lower model fitness and higher AUC (lower drug sensitivity). C-D. Drug target prediction accuracy (C) and signature target enrichment (D) of different signatures of PRISM drugs. Drugs with top and bottom model fitness were shown separately.

We first evaluated whether a reliable gene essentiality signature could be determined from the PRISM drug sensitivity profiles. Comparing with the CTRP and GDSC datasets, we observed a lower regression model fitness in the PRISM dataset (Dunnett’s test, all P-values < 0.001, **Supplementary Table 11**). As shown in **Figure 5B**, noncancer drugs tended to have lower drug sensitivities (median AUC of 1.1 compared to 0.71 and 0.90 for chemotherapeutic and targeted drugs, Dunnett’s test P-value = 1.6×10^-15^ and 8.9×10^-16^), which was associated with lower model fitness (Spearman correlation = −0.46, P-value < 2.2×10^-16^). In addition, compared to CTRP and GDSC, we observed that the model fitness decreased, even for the same set of cancer drugs (n = 44, **Supplementary Table 12**). Moreover, the gene essentiality signatures derived from PRISM were less consistent with those derived from CTRP and GDSC (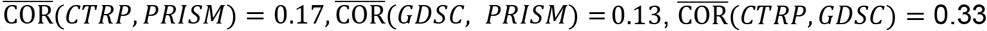, Dunnett’s test P-value = 9.4×10^-8^ and 1.6×10^-9^ for CTRP- PRISM and GDSC-PRISM, respectively).

We next evaluated the target prediction accuracy of the gene essentiality signatures, using the existing target annotation of the PRISM drugs as the ground truth (**Figure 5C** and **Supplementary Table 13**). We observed similar performance of the different signatures in predicting the targets (**Figure 5C,** left panel, n = 805). However, when we evaluated only the top 200 drugs with higher model fitness, we observed more significant improvement of the gene essentiality signatures (**Figure 5C**, middle panel, n = 200). Likewise, the quality of the gene essentiality signature associated with the target enrichment of the signatures. As shown in **Figure 5D**, we observed that the gene essentiality signature outperformed the gene expression signatures only for drugs with top model fitness (median AUROC of 0.72 versus 0.61, t-test P-value = 1.7×10^-3^). Together our results support the analysis of gene essentiality signatures only for PRISM drugs with top model fitness.

### Applying gene essentiality signature to explore mechanism of noncancer drugs

To minimize false predictions for the PRISM drugs, we considered the 312 drugs with top model fitness with a Spearman COR > 0.20 (**Supplementary Figure 6**). Furthermore, only the drugs with target information were retained, resulting in 222 cancer and 46 noncancer drugs. We found that the target prediction accuracy is much higher for cancer drugs than noncancer drugs (median AUC 0.97 and 0.26, t-test P-value = 3.8×10^-7^, **Figure 6A**). Likewise, the targets are better enriched in gene essentiality signatures of cancer drugs (median AUC 0.73 vs 0.64, two-sided t-test P-value = 0.04). We also observed that a noncancer drug’s putative targets were less likely to be shared with other drugs. **(Figure 6B)**. As shown in **Figure 6C**, target prediction was significantly improved for drugs having a target(s) shared by drugs in the training set (Dunnett test, all P-values < 0.001). There might be two explanations for such results. First, given that the putative targets of noncancer drugs indeed account for the drug’s killing effect, the low frequencies of the target genes in the training set prevented efficient model training. Secondly, using putative targets as the ground truth may lead to an underestimation of the prediction accuracy, especially for noncancer drugs, as cancer is not the targeted disease in the original development of noncancer drugs. The putative targets alone, therefore, may be insufficient in explaining the growth inhibition effect of noncancer drugs.

**Figure 6.**
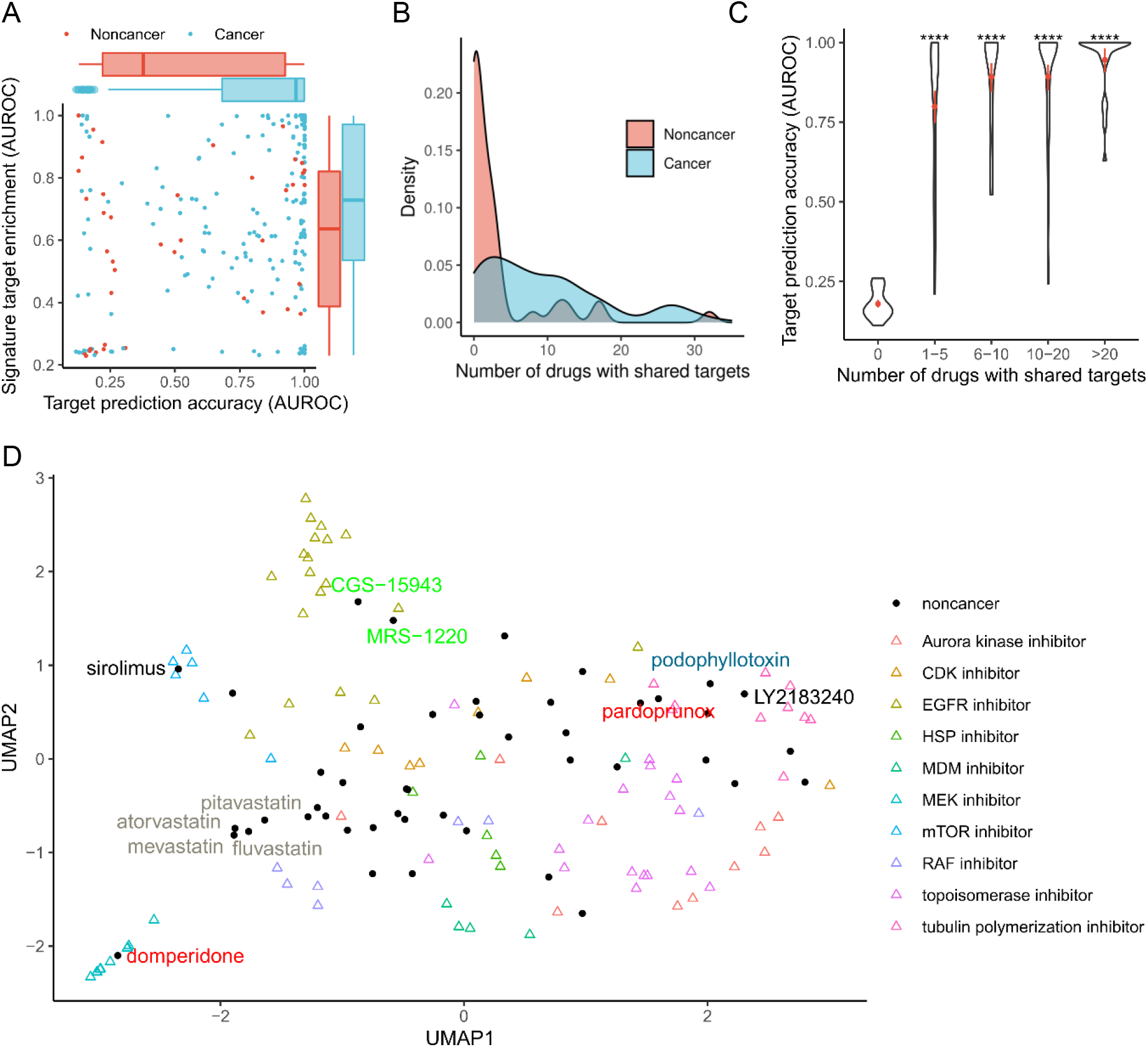
Target identification for noncancer drugs. A. Target prediction and gene signature enrichment of the putative targets of the PRISM drugs. B. Kernel density estimation of drugs having shared targets with other drugs C. Target prediction accuracy for drugs with different numbers of shared targets. D. UMAP projection of PRISM drugs by their gene essentiality signatures.

Given that the putative targets for noncancer drugs often cannot explain the mechanisms of action in cancer treatment, we further explored the similarity of gene essentiality signatures between cancer and noncancer drugs. We projected the gene essentiality signatures using the Uniform Manifold Approximation and Projection (UMAP). As shown in **Figure 6D**, cancer drugs with common mechanisms mostly clustered together, consistent with the clustering results determined by their drug sensitivity profiles^10^. Moreover, the gene essentiality signatures recovered putative anticancer mechanisms for multiple noncancer drugs. For example, sirolimus, an immune suppressor initially approved for preventing renal translation rejection, formed a cluster with mTOR inhibitors. Likewise, the anti-viral drug podophyllotoxin, later known to inhibit human tubulin polymerization, was clustered into cancer drugs as tubulin polymerization inhibitors. Detecting these clusters of well-studied cancer and noncancer drugs supports the validity of using gene essentiality signatures to recover putative mechanisms for non-cancer drugs for cancer treatment.

Furthermore, we observed novel proximity of a few noncancer drugs. For example, a FAAH inhibitor LY2183240 fell into the tubulin polymerization inhibitor cluster. Adrenergic receptor antagonists (CGS-15943 and MRS-1220) were found near the cluster of EGFR inhibitors. Interestingly, dopamine receptor antagonists were more diverse, with domperidone clustering with mTOR inhibitors, whereas pardoprunox clusters with tubulin polymerization inhibitors. Four HMGCR inhibitors (pitavastation, atorvastatin, mevastatin and fluvastatin) formed an isolated cluster that is clearly separated from the other drugs, suggesting their unique mode-of-action in cancer treatment. Together these results showed the capability of gene essentiality signatures to explore the mechanisms of noncancer drugs to rationalize drug repurposing.

### Experimental validation of noncancer drug target predictions

We applied the PISA assay to deconvolute the targets of noncancer drugs experimentally based on the solubility shifts. The PISA assay is a mass spectrometry-based technique increasingly used to identify protein drug targets at the proteome scale, and thus allows elucidating the mechanism of action in an unbiased manner^8,9,34,35^. We used Kuramochi ovarian cancer cell line as the model for the validation experiments.

A total of 20 noncancer drugs that showed sensitivity to Kuramochi were selected based on the PRISM and our in-house confirmatory screen data (AUC < 0.8, **Supplementary Table 14**). For these 20 drugs, we evaluated whether the predicted drug targets can capture the interacting targets identified by PISA (**Supplementary Table 15**). For each drug, we compared the protein soluble amount of drug-treated samples compared to DMSO controls in triplicate. The PISA scores were calculated by considering both fold change and statistical significance (see Method section for more details). To control the false positives, we evaluated only the targets that were shared between model predictions and top 25% PISA targets (median candidate target size n = 147), using Pearson correlation as the metric. We further generated 10000 random predictions to form a null distribution of the Pearson correlations.

We found that the average correlation for the 20 drugs was significantly higher than random prediction (**Figure 7A**, Pearson correlation 0.082 vs 0, two-sided P-value 0.024). The top performance was observed for the drug podophyllotoxin (COR 0.68, FDR < 0.01). We further identified the other four drugs all with FDR < 0.2, including harringtonine, fluvastation, pitvastatin, and lanatoside-c. Capable of stopping replication of both cellular and viral DNA, podophyllotoxin has been widely prescribed for genital warts and molluscum contagiosum. More recently, podophyllotoxin has been studied as an anti-tumor agent, being reported to destabilizes microtubules and prevents cell division^36^. Notably, the cancer target TUBB, together with several other tubulin family genes, were ranked at the top by both PISA experiments and our gene essentiality signature-based prediction (**Figure 7B**). Similar results were also observed for the two statin family drugs fluvastatin and pitvastatin, where HMGCR is confirmed by both PISA and our prediction for these two HMGCR inhibitors. Together our PISA experiment validated the target prediction of noncancer drugs.

**Figure 7.**
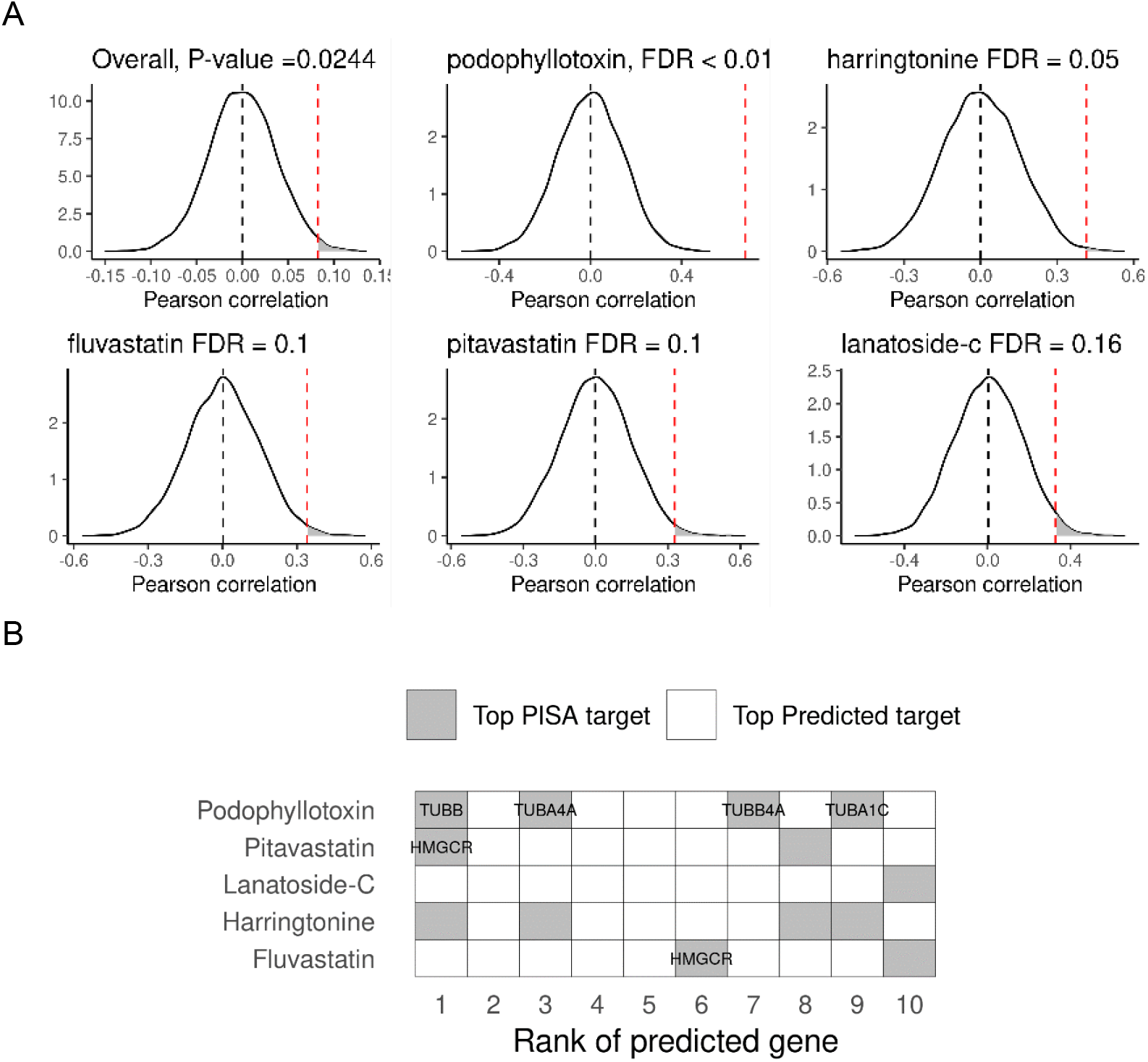
Experimental validation of predicted targets for noncancer drugs. A. Significance of the correlation between PISA scores and predicted scores. Gene essentiality-based prediction performance is marked as red dash line, while random predictions from 10,000 iterations formed a null distribution. B. Top PISA target genes are prioritized in predicted targets.

## Discussion

Characterizing the full spectrum of drug targets is fundamental for drug development^37^. In this study, we developed a computational method to determine gene essentiality signatures for drugs. Using the GDSC and CTRP datasets, we showed that our gene essentiality signatures outperformed gene expression signatures and conventional chemical fingerprints, in both supervised target prediction and *de novo* drug mechanism exploration. The gene essentiality signature enables drug-specific pathway analysis, helping uncover the mechanisms of action beyond primary targets. We further determined the gene essentiality signatures for drugs that were tested in the PRISM study^33^. We showed the feasibility of our method to predict the drug targets. More importantly, we predicted the cancer targets and their mechanisms of action for noncancer drugs.

We have applied the gene essentiality signatures to study cancer relevant targets for noncancer drugs. We have successfully validated the predicted target profiles for multiple drugs using PISA, a cell-based, proteome-wide target deconvolution assay. However, our validation results were based on a single cell line. To what extend the result can be generalized requires further investigation. On the other hand, the gene essentiality signatures were established with pan-cancer functional screening data, the predicted targets are hence less likely to be biased to certain type of cancer. Drug targets prioritized by both our prediction and PISA validations are more likely to be functionally relevant, providing a strong basis for further target validation studies.

To evaluate the performance of our prediction method, we leveraged existing knowledge about the drug targets as the ground truth. However, such information is likely to be incomplete, especially when evaluating noncancer drugs, as the primary targets previously identified for noncancer drugs may be irrelevant to their cancer-killing efficacy. Indeed, for noncancer drugs, we observed lower concordance between predicted and putative labels. It should be noted that for cancer drugs, focusing on primary targets still induce a risk of missing additional targets that account for the treatment effecacy^1,38^. In the future, we expect that the proteome-wide drug target deconvolution experiments could generate more unbiased data to improve computational methods for drug target prediction^8,39^.

Meanwhile, through our analysis of gene essentiality signatures of CTRP and GDSC drugs, we identified drugs that block DNA repair, which may help improve the efficacy of immune checkpoint inhibitors^31^. Immune checkpoint inhibitors have become standard-of-care in the first-line treatment of multiple cancers^40,41^. Despite their popularity, the responsiveness to immune checkpoint inhibitors varies greatly across individuals^42^. For example, we and others have identified that higher tumor mutation burden (TMB) serves as an important prognosis biomarker for immune checkpoint inhibitor treatment^43,44^. Using the gene essentiality signatures, we identified multiple compounds that may inhibit the DNA mismatch repair process of tumor cells (**Table 1**). Indeed, we have successfully predicted all the approved drugs that are under phase 2 clinical trials in combination with immune checkpoint inhibitors. Therefore, our gene essentiality signatures may support drug combination discovery to circumvent the limitation of immunotherapy.

Our method may be extended to study the mechanisms of action of drug combinations, as a large amount of drug combination screening data is also available for the modelling^45–47^. Secondly, we assumed that a drug inhibits its targets. However, drug target interaction may not always lead to an inhibition but rather a stabilizing or activating effect^48^. Such an assumption may contribute to false negative drug target predictions. In addition, we relied on the drug sensitivity data and gene essentiality data on cancer cell lines, which may not capture the mechanisms of action of drugs in the tumor microenvironment. Drug screening based on emerging in-vitro models, such as organoids and 3D cell culture^49,50^, may further rationalize drug discovery and drug repositioning^51,52^.

## Methods

### Gene essentiality data

We used the DepMap data portal^53^ to collect the gene essentiality profiles. Specifically, we collected the CERES gene essentiality scores derived from CRISPR screens^20^ and the DEMETER2 gene essentiality scores derived from shRNA screens^19^. Furthermore, we determined Combined Essentiality Score (CES^54^) by integrating CERES and DEMETER2 with the molecular profiles of the cell lines, including the TPM gene expression values derived from RNA-seq, the gene expression values from the microarray screens, the genelevel copy numbers, and the somatic mutations. We considered the three variants of gene essentiality features (CERES, DEMETER2, and CES) in the subsequent analyses. Genes with a missing rate of more than 20% were removed, after which the R MissForest package was used to impute the rest of the genes with the default parameters.

### Drug sensitivity data

We retrieved the CTRPv2 dataset from the NCI’s CTD-squared data portal, containing 545 small molecules and 907 cancer cell lines^55^. Cell viability percentages were determined at multiple doses of the drugs, after which a logistic curve was fitted. The area under the curve (AUC) was considered the drug sensitivity score, where an AUC of 0 represents complete cell growth inhibition, and an AUC of 15 represents no effect.

We also extracted the GDSCv2 data consisting of 198 small molecules across a panel of 809 cancer cell lines^56^. Furthermore, we obtained the PRISM data containing the drug sensitivity scores for 1,448 compounds against 499 cell lines^10^. In GDSC and PRISM, the AUC was calculated from the fitted dose viability curve and normalized further to the [0, 1] interval.

### Chemical fingerprints and gene expression signatures

The Python RDkit was used to generate Extended Connectivity FingerPrint (ECFP, 1024 features) and Molecular ACCess System fingerprints (MACCS, 256 features) for each compound based on its SMILES. In addition, we retrieved the LINCS L1000 Connectivity Map data to get the gene expression signatures^12^. The consensus drug signatures with 978 features were used following the procedures described in the previous work^57^.

### Drug target ground truth data

Binary drug targets were extracted from the corresponding drug screening studies, where either original drug library annotation or Drug Repurposing Hub^58^ (DRH) was used as the source. Furthermore, DrugTargetProfiler^59^ was employed to extract and integrate quantitative drug target data from DrugTargetCommons^60^, where measurements across multiple assays were harmonized into a consensus score between 0 (non-interaction) and 1 (strong interaction).

### Determination of gene essentiality signatures

We built a drug sensitivity prediction model using three variants of gene essentiality signatures (CES, CERES, and DEMETER2). For each *drug_i_*we fit an L1-penalized multivariate linear regression model *M_i_* by regressing its drug sensitivity AUC against genome-wide gene essentiality for multiple cell lines:

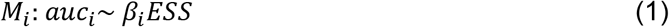

Where *auc_i_*, is a 1 × *n* vector, and *ESS* is an *m* × *n* matrix representing the essentiality profiles of m genes for the n matching cells. The 1 × *m* coefficient vector *β_i_* is then extracted to represent the gene essentiality signature of the drug.

Nested cross-validation (CV) was used for evaluating the model fitness, where the hyperparameters were tuned with a ten-bootstrap inner layer and the prediction accuracy was evaluated on the outer layer of five folds with three replicates. Model fitness was estimated using the holdout sets from the CV, including accuracy metric, i.e., coefficient of determination (R^2^), Spearman correlation coefficient (Spearman COR), area under the precision-recall curve (PRAUC), area under the receiver-operator curve (AUROC), as well as error metrics, i.e., mean absolute error (MAE) and root-mean-square error (RMSE). Regression metrics were calculated using the original numeric drug sensitivity scores and classification metrics were calculated by binarizing using the 90% percentile as the drug sensitivity threshold.

The overall difference among the three variants of gene essentiality signatures was evaluated by ANOVA test, and if significant, followed by post hoc Tukey’s test or Dunnett’s test. For each drug, the performance of the three variants of gene essentiality signatures was compared using paired t-test with Bonferroni correction.

### Evaluation of gene essentiality signatures

We evaluated which drug features (i.e., gene essentiality signatures, gene expression signatures, and chemical fingerprints) predicts drug targets. We considered a drug-target prediction method that was developed in recent competitions, significantly outperforming the other methods in an independent experimental validation^15^. This method is conceptually similar to the *k* nearest neighbors learning, where the targets of a queried drug are predicted by their neighbors. For a queried *drug_i_*, we determined its correlations with the training drugs in the feature space as shown below:

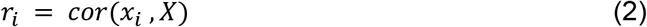

where represents the feature vector for the queried drug, *X* is a *m* × *n* feature matrix for the n training drugs. The drug targets for drug i were then predicted as an average of the drug target score of the training drugs, weighted by the correlation coefficients transformed by rectified linear activation function (ReLU):

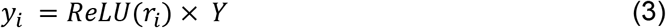

Where the *p* × *n* matrix, *Y* represents the target profiles for all the n training drugs in p genes, *ReLU*(*r_i_*) is the transformed coefficient vector and *y_i_* is the predicted target profiles for *drug_i_*. Cross validation with five folds and three replicates was employed to estimate the AUROC against the ground truth.

In addition, we evaluated the similarity of a drug pair based on multiple feature spaces. For each drug pair, we used Spearman correlation to determine the similarity in the drug sensitivity space, as well as in the predicted drug target space. We used the Tanimoto distance to determine the similarity in the ground truth target space, as well as in the chemical fingerprint space. We considered pairs of drugs with similar drug sensitivity profiles and drug-target profiles to be mutually repurposable. For example, for drug pair A and B in the CTRP dataset, we considered them mutually repurposable if both their drug sensitivity and drug target similarity are higher than 95% of other drugs paired with drug A or drug B. The threshold was set to 90% in GDSC due to insufficient ground truth drug pairs. For each mutually repurposable drug pair, we checked whether their similarity in the predicted target space is also at the top. The performance of different drug signatures was compared using the paired Wilcox signed rank test with Holm’s correction for multiple testing. In addition, only the top mutually repurposable drug pairs in the CTRP dataset (N = 16, with drug sensitivity similarity > 99%, target similarity 100%, and identical mechanisms of action) were shown.

### PPI network and pathway analysis

To study the functional relevance of the gene essentiality signatures, we mapped the genes in the PPI network constructed in the recent publication^24^. Each signature gene was annotated as the drug target, first-degree, second-degree, or other neighbors, depending on the shortest path between the gene and the putative target of the given drug. GSEA^61^ (Gene Set Enrichment Analysis) was conducted for the gene essentiality signatures against the KEGG and GO pathway gene sets collected from the MSigDB database^62^. Pathway associations were filtered with absolute normalized enrichment score (NES) > 1.5 and P_adj_ < 0.05.

### Cell culture

Kuramochi (JCRB Cell Bank, JCRB0098) cells were grown in RPMI 1640 medium (Gibco) supplied with 10% fetal bovine serum (Gibco), 1% GlutaMAX (Gibco) and 1% Penicillin-Streptomycin (10,000U/ml, Thermo Fisher Scientific). The passage and freezing steps were followed according to the handling procedure from ATCC.

### Drug treatment and sensitivity fitting

To calculate the dose-response curves for selected drugs, the drug treatment was designed and performed using the D300e dispenser (Tecan). Kuramochi cells were seeded at the density of 1000 cells per well in a 384 well black clear bottom microplate (Corning), on the same day, a stock of each drug was loaded to D300e dispenser and drugs were automatically loaded to each well according to the plan pre-made through D300e control software (Tecan). Each drug was tested for 10 concentrations in triplicate, starting from 10umol to 1nmol. At the desired end point, cell viability in individual well was measured using CellTiter-Glo® Luminescent Assay (Promega). Percentage inhibition was calculated using the following formula, 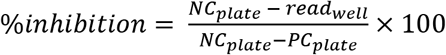, where NC stands for negative control (DMSO and MQ with tween for DMSO soluble and water soluble compounds, respectively) readout and PC stands for positive control (Benzoyl chloride) readout. Adding tween to water is required for dispensing water-based compounds using D300e dispenser. Dose response curves were then fitted using 4 parameter logistic function in drc R package.

### PISA assay

Kuramochi cells were cultured and treated for one hour with each compound, using a biological triplicate of each drug treatment plus DMSO control, for a total of five drugs per PISA assay. A total of four PISA assays in the 18-plex format were used for the 20 drugs, according to the following scheme for each PISA assay: 1) albendazole, AZ3146, BMS- 387032, digoxin, EVP4593; 2) fluvastatin, harringtonine, JIB04, lanatoside-C, monensin; 3) nanchangmycin, narasin, NSC-697923, oligomycin-A, parbendazole; 4) pitavastatin, podophyllotoxin, proscillaridin-A, puromycin, thiostrepton.

PISA was carried according to the published protocol^8^ optimized for higher proteome depth and 18-plex multiplicity using Tandem Mass Tag (TMT) pro (Thermo Fisher). Sample processing for proteomics based on nanoscale liquid chromatography and tandem mass spectrometry (nLC-MS/MS) was performed according to the PISA assay described for 18- plex format^9^ using a gradient for thermal denaturation as described previously^34^. Briefly, 50 μg of the soluble fraction of each sample after ultracentrifugation was reduced, alkylated, proteins precipitated and then digested using first LysC and then trypsin. Digested samples were then labelled with TMTpro 18-plex reagent kit. The final multiplex peptide sample was cleaned and desalted and fractionated by reversed phase at high pH into 48 fractions using a capillary off-line HPLC. nLC-MS/MS analysis was performed on all fractions using 95minute gradient per run (total run time 120minutes per sample) with a system composed by a nano-LC Dyonex connected to a Orbitrap Exploris™ 480 mass spectrometer (Thermo Scientific).

Analysis of nLC-MS fractions was carried out using Proteome Discoverer 2.5 (Thermo Scientific) with database search against the full Uniprot human protein database UP000005640 for peptide and protein identification and quantification. Datasets were normalized on total TMT ion reporter intensity and on the average of DMSO controls, to obtain the final values soluble amount variation of each protein and relative statistical significance of variation by P-values calculated by student’s t-test across their biological triplicates.

### Statistical analysis

All statistical tests were performed as two-sided tests with R (v4.0.5). Nonparametric tests were used to compare the rank and the parametric tests were used to compare the actual numerical values. Wilcox signed rank test was used for groupwise rank comparisons. Fisher’s test was used to compare the frequencies and t-test was used for two group numerical comparisons. Tukey’s test was used for multigroup pairwise post-hoc comparisons and Dunnett test was used to compare each group with the reference. Specific test statistics, degree of freedom, confidence interval and significance to reject the null hypothesis were reported in the supplementary tables. The standard error and confidence interval for mean estimates were shown as black dot and error bar in the violin plots. The marginal boxplots described the median (middle line), 25% and 75 quantiles (lower and upper hinges) as well as 1.5 multiples of interquartile range (whiskers).

## Supplementary Info

**Supplementary Figure 1.**
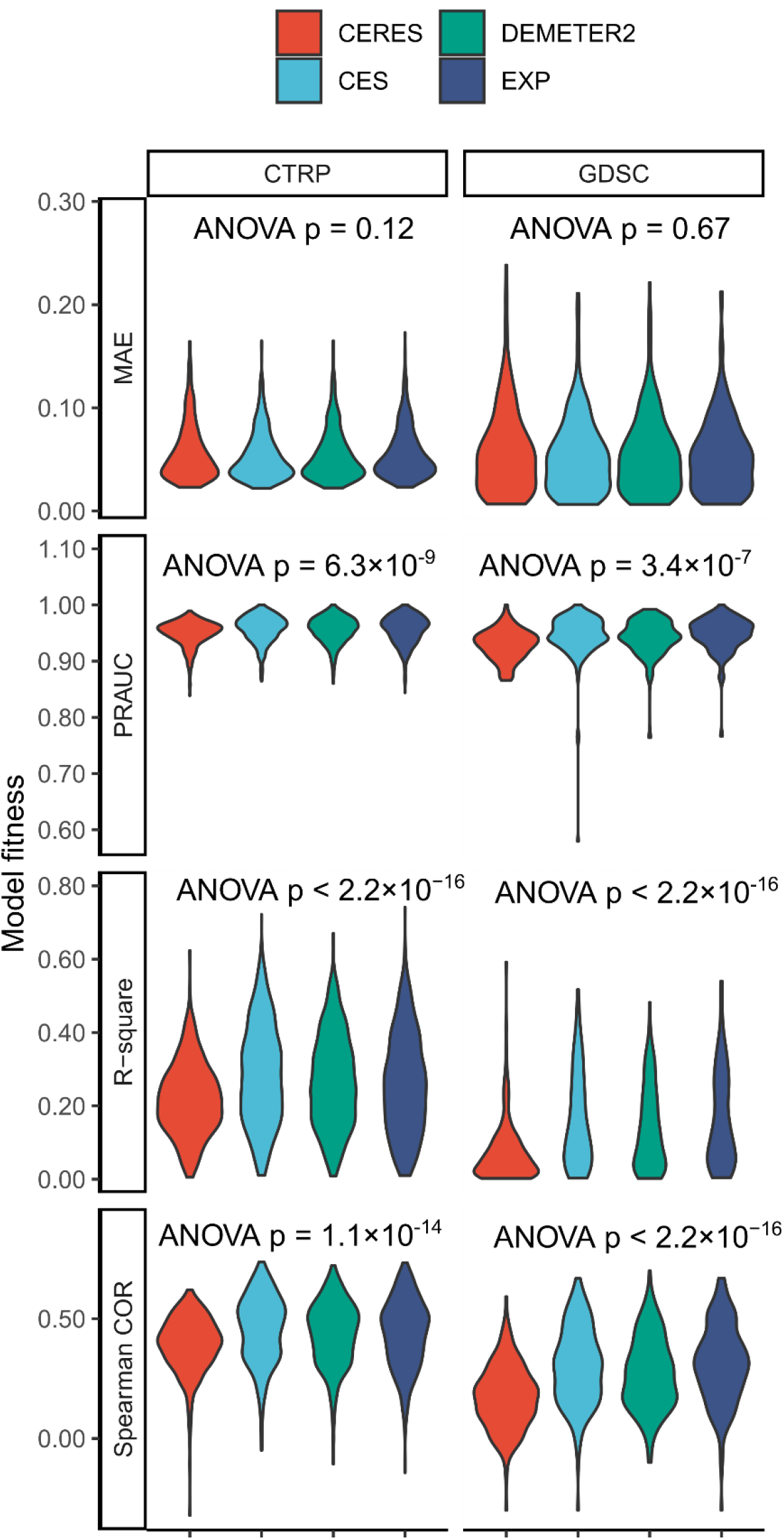
Model fitness of different gene features using cross-validation.

**Supplementary Figure 2.**
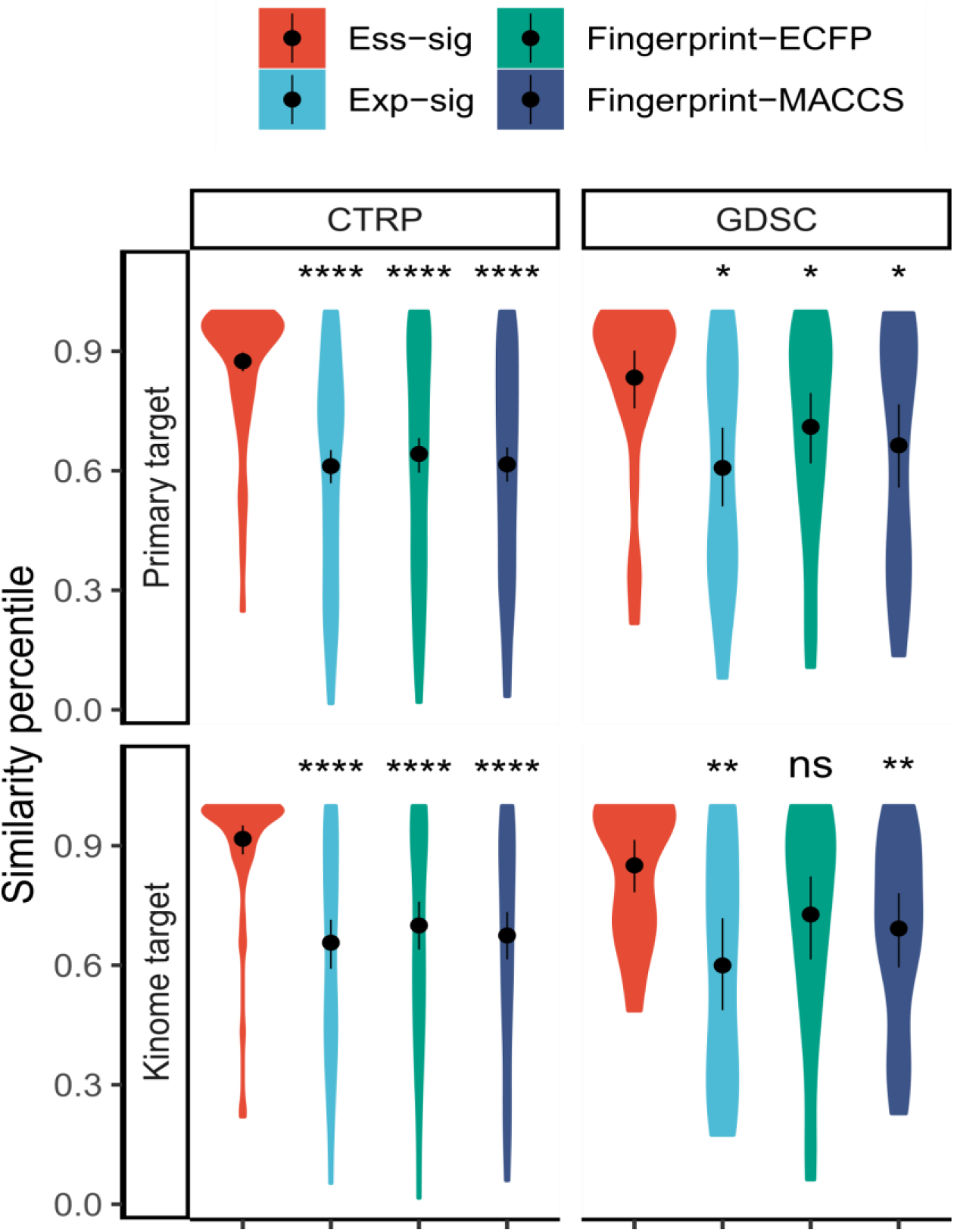
Rank of similarity for repurposable drug pairs. ns: p > 0.05, *: p <= 0.05, **: p <= 0.01, ***: p <= 0.001, ****: p <= 0.0001 by the paired Wilcox signed rank test with the gene essentiality signature as the reference. Ess-sig: gene essentiality signature, Exp-sig: gene expression signature.

**Supplementary Figure 3.**
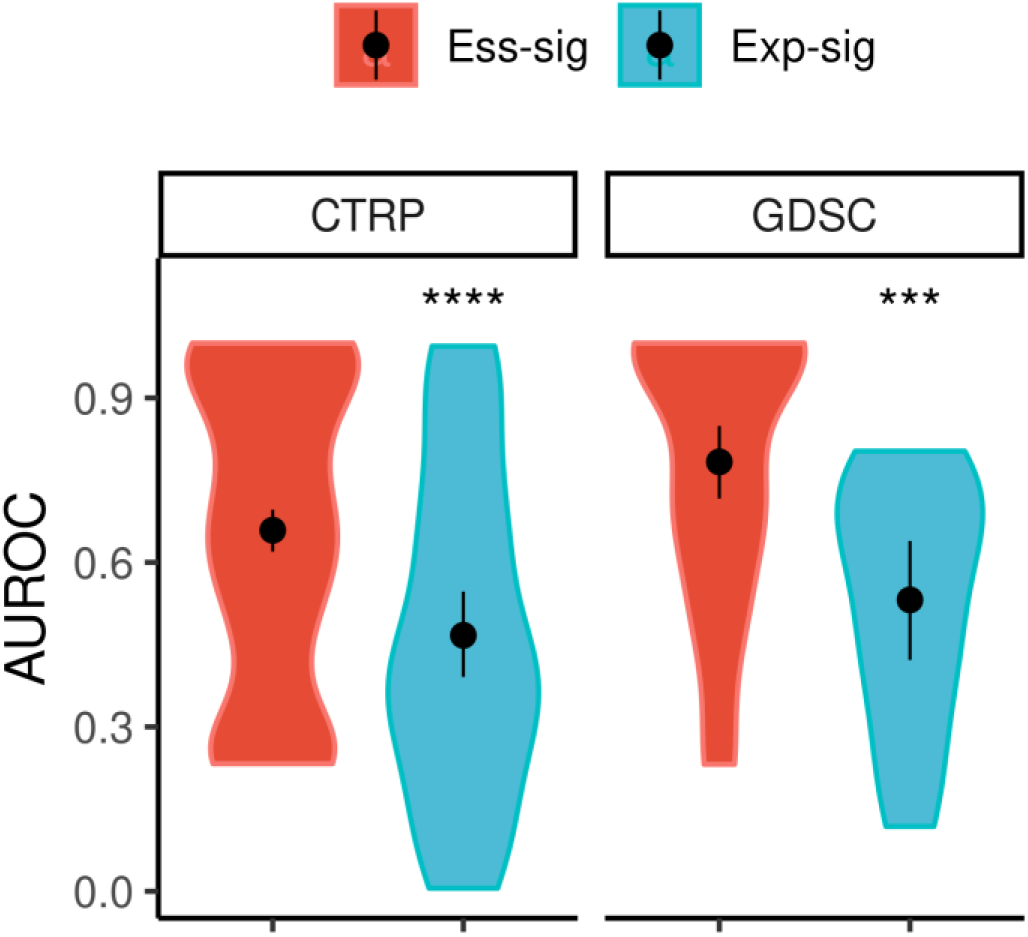
Accuracy of *de novo* prediction of primary targets.

**Supplementary Figure 4.**
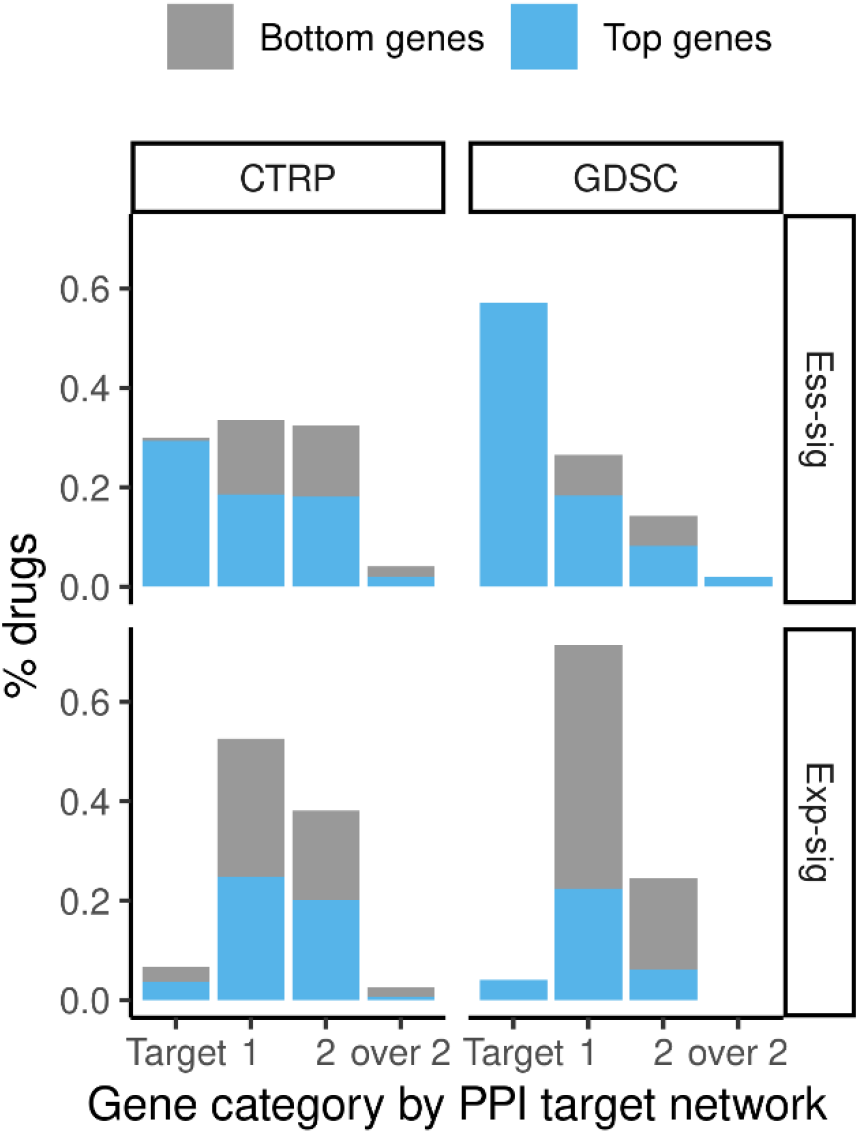
Percentage of drugs the target neighbors of which are prioritized by their top signature genes (top/bottom 50 genes).

**Supplementary Figure 5.**
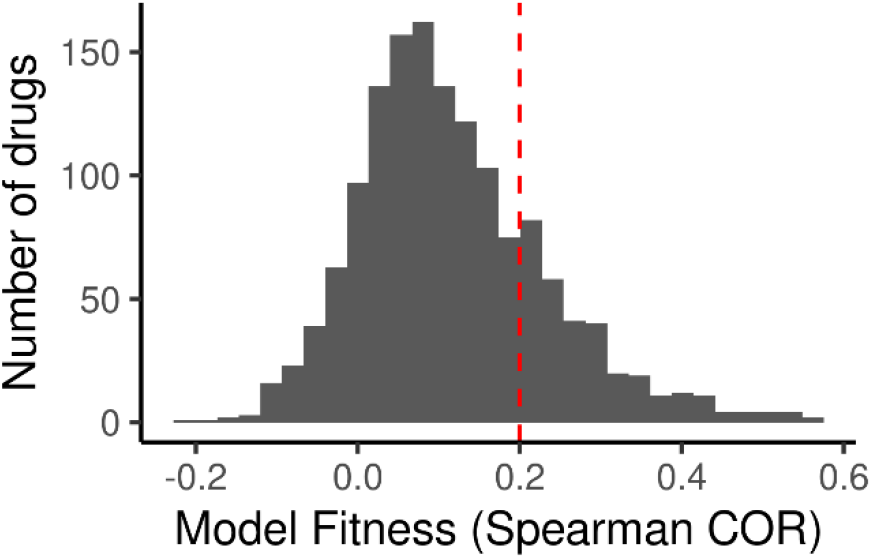
Distribution of model fitness for PRISM drugs.

**Supplementary Table 1** Mean estimates of drug sensitivity model fitness.

**Supplementary Table 2** Comparison of drug sensitivity model fitness between CES and CERES.

**Supplementary Table 3** Comparison of drug sensitivity model fitness between DEMETER2 and CERES.

**Supplementary Table 4** Comparing variance of RMSE for drug sensitivity model fitness.

**Supplementary Table 5** Statistical significance for comparing the accuracies in target prediction.

**Supplementary Table 6** Statistical significance for comparing the similarity percentile of repurposable drug pairs.

**Supplementary Table 7** List of KEGG pathways associated with the drugs’ gene essentiality signatures.

**Supplementary Table 8** List of Gene Ontologies (GO) terms associated with the drugs’ gene essentiality signatures.

**Supplementary Table 9.** Immuno-oncology (IO) combination clinical trials involving drugs associated with DNA mismatch repair.

**Supplementary Table 10** PRISM drug target annotation.

**Supplementary Table 11** Gene essentiality model fitness for CTRP, GDSC, and PRISM drugs.

**Supplementary Table 12** Gene essentiality signatures for the drugs screened in all the three studies.

**Supplementary Table 13** Performance of target prediction for drugs with varying gene essentiality model fitness.

**Supplementary Table 14** Drug sensitivity validation of noncancer drugs.

**Supplementary Table 15** PISA drug target deconvolution experiments of noncancer drugs.

## Data availability

Supplementary tables are available online. The datasets are publicly available in the following repositories, CTRP [https://portals.broadinstitute.org/ctrp.v2.1/], GDSC [https://www.cancerrxgene.org/] and DepMap [https://depmap.org/portal/]. DTC [https://drugtargetcommons.fimm.fi], DTP [http://drugtargetprofiler.fimm.fi/], Drug Repurposing Hub [https://clue.io/repurposing#download-data]. Specifically, DepMap 21Q1 release was employed to derive the input datasets including, CERES (Achilles_gene_effect.csv), TPM RNA-seq gene expression matrix (CCLE_expression.csv), the gene level CNV file (CCLE_gene_cn.csv), and the mutations (CCLE_mutations.csv). In addition, DEMETER2 (D2_combined_gene_dep_scores.csv) was collected from DEMETER2 Data v6. The gene expression from the microarray (CCLE_Expression_Entrez_2012-09-29.gct) was collected from the CCLE release available in DepMap portal. For the CTRP dataset, the curve-fitting parameters from the post-quality control dataset (v20). For GDSC we employed the curve statistics (GDSC2_fitted_dose_response_25Feb20.xlsx) from its version2. PRISM data (secondary-screen-dose-response-curve-parameters.csv) was collected from its 19Q4 release.

## Code availability

The GitHub repository is available at https://github.com/Wenyu1024/drug_moa.

## Acknowledgment

We appreciate the open science data sharing of the Broad Institute of MIT and Harvard for providing The Cancer Dependency Map (DepMap), PRISM, and the Drug Repurposing Hub. We thank CSC, Finland for providing the IT services. The illustrative cartoons in Figure 1, Figure 2A, and Figure 4D were generated using bioRender. The Chemical Proteomics core facility at Biomedicum (MBB, Karolinska Institute), also Unit of SciLifeLab and unique Chemical Proteomics node of the Swedish National Infrastructure for Biological Mass Spectrometry (BioMS), provided full support in the experimental design and performance of the proteomics using the Proteome Integral Solubility Alteration (PISA) assay.

## Author contribution

The work was supervised by J.T. W.W. and J.T. conceived of the study. W.W. developed the models and led the computational analysis. S.Z., Z.T., Y.W., and J.A. contributed to the data collection, while J.B., S.H, and J.E. contributed to the interpretation of the findings. J.B conducted drug sensitivity experiments. XZ and MG designed and conducted the PISA experiments. W.W and J.T. wrote the manuscript. All authors have read and agreed to the final version of the manuscript.

## Corresponding authors

Correspondence to Wenyu Wang or Jing Tang.

## Funding

This work was supported by the EU H2020 (EOSC-LIFE, No. 824087, JT), the European Research Council (DrugComb, No. 716063, JT), and the Academy of Finland (No. 317680, JT). W.W. was funded by the FIMM-EMBL international PhD program, Doctoral Program of Biomedicine at University of Helsinki, Cancer Foundation Finland, K. Albin Johanssons stiftelse, Ida Montinin Säätiö, Orion Research Foundation sr. and Biomedicum Helsinki Foundation. S.H was supported by the National Natural Science Foundation of China (No. 82203120) as well as the Key Research and Development Program of Shaanxi (Program No.2022SF-092).

## Competing interests

None to declare

## Notes

### Competing Interest Statement

The authors have declared no competing interest.

### Summary of Updates

We have extended the manuscript with an additional section describing experimental validation.

